# Stopping and Changing Expected and Unexpected Movements

**DOI:** 10.64898/2026.02.16.706101

**Authors:** Simon Weber, Nicholas Vucak, Sauro E. Salomoni, Alison J. Ross, Emily Coleman, Mark R. Hinder

## Abstract

The capacity to cancel or adapt planned actions in response to changing environmental demands is essential for navigating our complex world. While past research has shown that an individual’s expectations of upcoming movement demands influence the speed of action initiation, the effect this has on subsequent cancellation or adaption of that movement remains unknown. 25 healthy adults completed stop signal tasks and stop change tasks in which biasing cues (e.g., “70% left”) accurately indicated the probability that a left, or right button press would be required. As expected, responses that were congruent with the cue were faster than incongruent responses; however, biasing cues had no effect on behavioural or physiological (electromyographical) indices of stopping speed.

Stopping latencies were found to be *faster* in the stop change task than the stop signal task, corroborating other recent work. However, a second experiment (25 healthy adults) which used the same stimuli for both tasks (varying only the instructions), revealed no difference – highlighting the sensitivity of the stop process to stimulus effects, and a common confound in the literature.

We also observed that physiological indices of action reprogramming (following a stop) were faster in congruent than incongruent trials. Collectively, these results suggests that preparatory changes that accompany expected movements influence the enaction of movement both prior to, and after stopping, but the stop mechanism itself, remains independent of these preparations. These results inform how action cancellation and adaption are applied in real world environments, where expectations continually interface with our motor plans.

**Highlights:**

* Anticipating a movement increases the speed of its enaction but not subsequent cancellation
* Expected movements can be reprogrammed more quickly than unexpected movements
* The latency of action cancellation is highly sensitive to stimulus effects

## Introduction

Central to flexible and goal-directed behaviour is the capacity to cancel or modify planned actions in response to environmental changes. This capacity, usually referred to as *response inhibition*, is widely researched in cognitive and behavioural neuroscience (Albaladejo-García et al., 2023; Lipszyc & Schachar, 2010; Verbruggen & Logan, 2008). The gold standard experiment to investigate this phenomenon in controlled laboratory environments is the stop signal task (SST; Lappin & Eriksen, 1966), which typically requires participants to perform a choice response (left- or right-hand) to a visual “go signal”. On a minority of trials, a subsequent “stop signal” occurs, requiring the inhibition of the planned or initiated response (Verbruggen et al., 2019). Manipulation of the delay between the go and stop stimuli allows estimation of “stop signal reaction time” (SSRT), an indirect measure of the speed with which someone can stop a movement. Stopping deficits (slower SSRTs than healthy controls) are observed in various clinical disorders (Lipszyc & Schachar; 2010) and are known to degrade as we age (Rey-Mermet & Gade, 2018).

Despite their proliferation, SSTs capture a very narrow range of the diverse real-world contexts in which we need to cancel, *or change*, our actions. At any given time, our actions (planned and initiated) reflect a combination of expectations and prior knowledge about our environment. For example, suppose we are anticipating a tennis serve from a player we know usually aims their serve to the left. If our prediction is correct, we can react to the serve more quickly, performing the requisite movements more efficiently than if our prediction had been incorrect (i.e., *incongruent* with prior expectations). While the psychological and neurophysiological processes are not fully understood, facilitation of movements has been well established in controlled laboratory settings using biasing cues (e.g., “70% left”) before choice responses (Miller & Low, 2001; Garton et al., 2019). In the current experiments we sought to determine what happens if *stopping* is required in this context. For instance, if the ball is served in the direction we expect, but we recognise it is going out of the court, does our prior expectation influence how quickly we can stop and adjust out planned return? At the time of writing, no published research has investigated whether explicitly biasing the *go* response influences the subsequent cancellation of movement, despite the extensive literature on how biasing influences movement enaction.

The independent race model (Logan, et al., 1984) that underpins SSRT calculation would predict that biasing of the go response should *not* influence the speed of the upcoming stop, as the stop and go processes are considered independent. However, past research has shown that both the discriminability (Ma and Yu, 2016), and sensory modality (Weber et al., 2024b) of the go stimulus can influence the influence the speed of stopping. Specifically, when go stimuli are hard to discriminate stopping speed is faster (Ma and Yu, 2016), and when go stimuli are presented in a different modality to the stop stimuli (e.g., visual go auditory stop, or vice-versa), stopping is faster (Weber et al., 2024b). Given these conflicting accounts, we designed novel SSTs that incorporated trial-level biasing cues providing probabilistic information regarding left or right responses, to see if this influenced subsequent stopping speed.

Despite the popularity of the SST, everyday situations rarely demand pure suppression of actions. Rather, they often require the abandonment of obsolete planned actions followed by the adaptation of behaviour. Addressing this complexity, Logan (1994) developed the stop change task (SCT) as an extension of the original SST. The SCT is similar to the SST, in that a subset of trials requires the inhibition of a planned/initiated action. However, the SCT also requires subjects to replace the stopped response with a *new* response, reflecting the behavioural adaptation demanded by everyday actions. Thus, change trials in the SCT involve three conceptually distinct stages: *go, stop*, and *re-engagement*. While the SCT has been used to identify inhibitory deficits in a range of clinical populations (Geurts et al., 2004; Bekker et al., 2005; Boecker et al., 2013), there is ongoing debate as to whether the inhibitory processes involved in the SCT are the same as those in the SST (Boecker et al., 2013). For instance, while some studies have observed longer stopping latencies in the SCT relative to the SST (Boecker et al., 2011; Logan & Burkell, 1986; De Jong et al., 1995; McClure et al., 2005), others have observed no difference (Drueke et al., 2010) or *faster* stopping in the SCTs than SSTs (Hervault & Wessel., 2025). Notably, most research comparing stopping speed in SST and SCTs has done so solely using SSRT calculations. The use of SSRT in SCTs requires the assumption that the stop and change responses are performed in serial, and that the additional movement after the stop does not interfere with go and stop processes (Verbruggen et al., 2008). Notably, SSRT calculations were conceived based on the standard SST (a simple race between a stop and go process) and may not generalise to complex stopping tasks in which additional (changed) responses are required (Gronau et al., 2024). One solution to this problem is to use electromyography (EMG) to enable direct measurements of the timing of initiation and cancellation of actions at the level of the muscle. Recording EMG data allows the observation of sub-threshold (i.e., too low to evoke a behavioural response) muscle activation on successful stop trials. These instances, referred to as *partial bursts*, occur when a motor response is initiated and subsequently cancelled before sufficient force is generated to press the button (i.e., a *covert response*). The delay between the stop or change signal and the peak amplitude of the partial burst (i.e. when muscle activity begins to decrease) is referred to as EMG *CancelTime* (Jana at al. 2020; Raud et al., 2022). CancelTime is considered a robust measure of stopping latency, as it provides a trial-level measure of inhibition, as opposed to SSRTs which only provides a single *estimate* of a person’s stopping speed over the whole experiment or all trials within a particular condition. Furthermore, CancelTime is well-suited to both SST and SCT contexts, avoiding the questions of validity regarding SSRT calculations in the more complex task. To the authors’ knowledge only one other study has compared CancelTime in SSTs and SCTs (Hervault & Wessel., 2025), observing faster stopping in change tasks then stop tasks, though the reason for this was unclear, and further investigation is justified. EMG also allows for a trial-level analysis of how quickly a movement is reprogrammed (that is, the time from when suppression occurs to when the generation of muscle activity begins on the other hand; Weber et al., 2024a).

In sum, the current experiment aimed to investigate the effect of biasing the go response on both stopping and changing, applying physiological measures that allow for direct observation of stopping and reprogramming speed at the level of the muscle.

## Experiment 1 Methodology

### Participants

Twenty-five healthy participants (13 female; M_Age_ = 31 years, SD = 9.8 years) were recruited either through the University of Tasmania psychology research participation scheme, or by direct invitation, for research credit or a $20AUD gift voucher, respectively. This research was approved by the Tasmanian Human Research Ethics Committee (reference number H0016981).

### Experimental Setup

Participants sat approximately 80cm from a computer screen with their forearms pronated (palms down) resting on a desk. Each hand was positioned on cushioned platforms so that their index fingers were adjacent to response buttons that were mounted in the vertical plane. To record a button press, participants performed horizontal abduction with the left or right index finger (based on the visual stimulus presented). Visual stimuli and behavioural responses were presented and captured using PsychoPy3 (Peirce et al., 2019).

EMG recordings were obtained using adhesive electrodes arranged in a belly-tendon montage on the first dorsal interosseus (FDI) muscle on each hand, with a grounding electrode on the head of the ulna. The FDI muscle was selected because it acts as a primary agonist during index finger abduction, as required to press the button. To maximise the activation of this muscle and improve signal-to-noise ratio (SNR), voluntary button presses were conducted using abduction (with buttons mounted vertically, as opposed to pressing ‘down’ on buttons or keys, where FDI only acts as a co-agonist resulting in a lower SNR). Notably, FDI abduction allows for cleaner EMG data compared to extrinsic finger flexors (the primary agonists if buttons were mounted horizontally) as there is less crosstalk from task-irrelevant forearm muscles, Salomoni et al., (2025).

To determine the correct placement of electrodes, participants performed an isometric abduction of their index finger to activate the FDI muscle. EMG data was recorded using Signal (Cambridge Electronic Design Ltd.), with recording synchronised to behavioural responses. EMG activity was recorded in sweeps beginning 500ms prior to stimulus presentation for a duration of 2000ms. The experimenter monitored the output and reminded participants to relax their hands if noise from unintended muscle activity was detected.

Prior to starting all tasks, participants were presented with comprehensive written instruction screens within PsychoPy3, as well as verbal confirmation from the experimenter.

### Task Overview

Participants completed a cued choice response task (CRT) in which they were instructed to respond with unimanual button presses as quickly as possible to the location of stimuli presented on screen. In all trials, a yellow (or cyan) circle appeared on either the left- or right-hand side of the screen (i.e., the *primary*, or *go* stimulus; see Figure 1a). Stimuli on the left of the screen required a left button press, and vice versa. The colours were counterbalanced such that some participants responded to yellow (N = 12) and some to cyan (N = 13) go stimuli. Prior to the go stimulus presentation, a biasing cue provided accurate information regarding the upcoming stimulus location in the form “70% likely left” or “70% likely right.” Thus, stimuli could be either congruent or incongruent with the preceding cue. Feedback was given after each trial. On successful trials, participants were shown their response time. Incorrect responses generated the feedback ‘Wrong response,” and missed trials (where no response was made), “Missed.”

**Figure 1:**
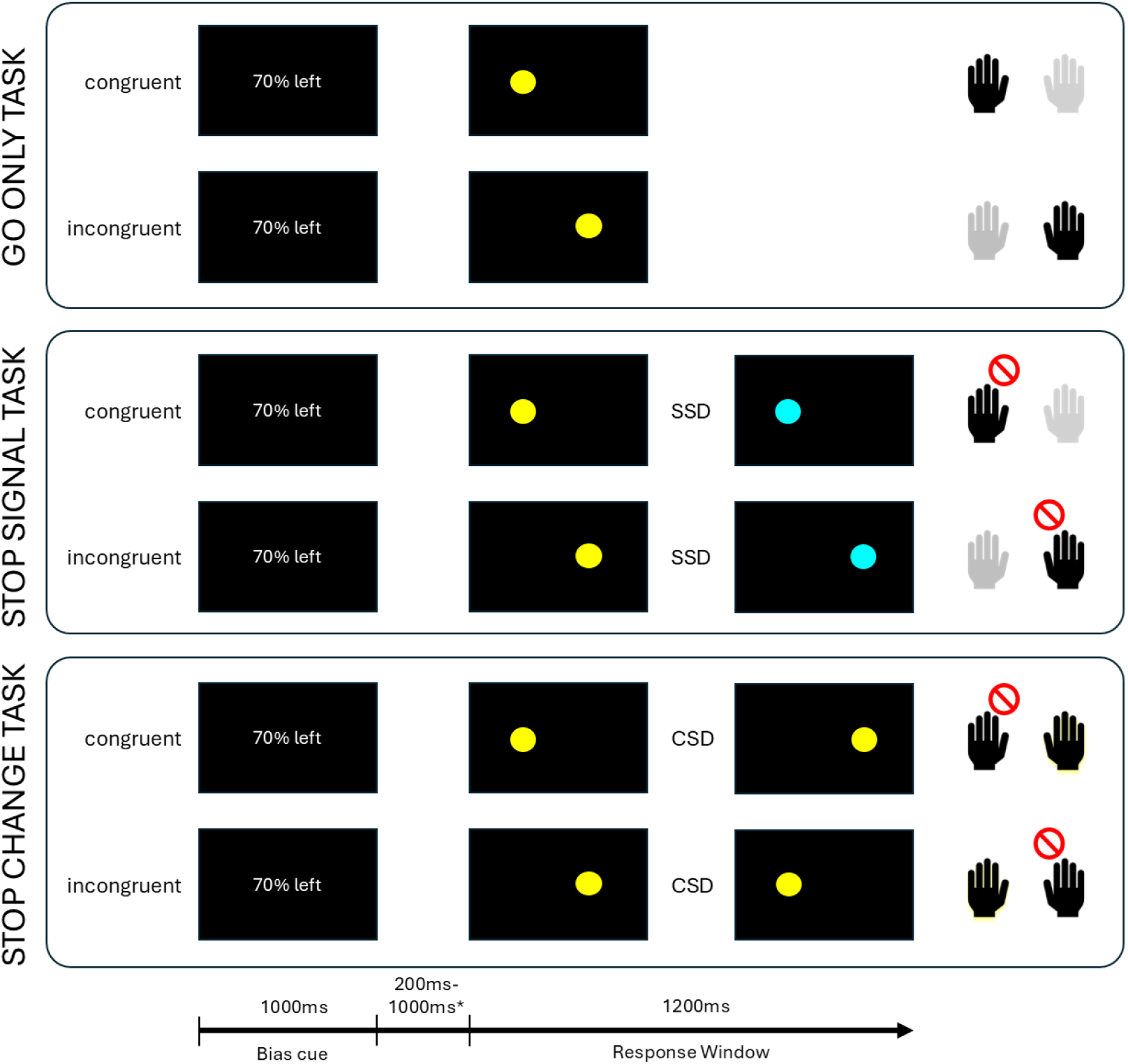
Timings and stimuli are depicted for the go-only task (top), the stop signal task (SST, middle), and the stop change task (SCT, bottom). The first column shows the biasing cue which accurately reflected the primary (go) stimulus (second column), while the third column represents the secondary stimulus (following an SSD or CSD delay, depending on task). For each trial type, correct responses for each hand are indicated in the fourth column: Grey hands are non-selected (were not cued by the go stimulus), red ‘stop’ markers indicate that the response should be cancelled (no button pressed). The change response is denoted as the hand outlined with yellow. Note that in the SST and SCT conditions, stop and change trials only constituted one-third of all trials; the remaining two-thirds of trials were go trials, as depicted in the top panel. **\***pre-stimulus delay time was randomly generated within a range of 200-1000ms from a truncated exponential distribution. SSD=Stop Signal Delay; CSD = Change Signal Delay.

Participants first completed 30 go-only CRT trials (referred to as the Go Only condition; see Figure 1). This was completed first to establish baseline RTs for both congruent and incongruent responses. They then progressed to the main phase of the experiment, where stopping was incorporated into the CRT under two conditions (conducted separately; counterbalanced): stop (the stop signal task, SST) and change (the stop change task, SCT). The colour of the go and stop stimuli were counterbalanced, such that for some participants, yellow served as go and cyan as stop (N = 13), and for others, cyan was go and yellow was stop (N = 12). The SST and SCT both involved 600 trials divided into 12 blocks of 50 trials with short user-defined breaks (at least 30 seconds) after each block to minimise fatigue. Overall, the experiment included 1230 trials and took approximately two hours to complete.

In the SST condition, two-thirds of trials (400 trials) were go trials (described above), while the remaining third (200 trials) were stop trials. During stop trials, the colour of the primary stimulus changed after the “stop signal delay” (SSD), requiring participants to inhibit their response (i.e., make no button press; see Figure 1). Both go and stop trials all followed accurate informative biasing cues regarding the location of the upcoming go stimulus, such that for go trials 280 were congruent and 120 were incongruent, while for stop trials 140 were congruent and 60 were incongruent. SSD was set at 150ms for the first stop trial and was adjusted following each subsequent stop trial, such that following a successful stop (no buttons pressed) SSD decreased by 50ms (making the next stop harder) and following a failed stop (either button pressed) SSD increased by 50ms (making the next stop easier; Verbruggen et al., 2019). Separate SSD staircasing procedures were used for congruent and incongruent stop trials (e.g., performance on congruent stop trials would not influence SSD for incongruent stop trials).

### Stop change Task (SCT)

In the SCT condition, two-thirds of trials (400 trials) were go trials, while the remaining third (200 trials), were change trials. In change trials, after a predetermined change signal delay (CSD), the primary stimulus disappeared from its original position and simultaneously re-appeared on the other side of the screen. The CSD was staircased in the same way as the SSD in the SST. In change trials, the participant was required to inhibit the initiated response to the original go stimulus and subsequently respond with the contralateral hand. For example, if a yellow circle appeared on the left and quickly changed to the right side of the screen, it required both the inhibition of the left-hand response and subsequent button press with the right-hand to register a correct response. As in the SST, accurate biasing information preceded all trials (for go trials 280 were congruent and 120 were incongruent, while for change trials 140 were congruent and 60 were incongruent). Feedback for the SCT was the same as the SST (no RTs were given for the response in change trials, instead the feedback simply read “correct”).

In both the stop and change conditions, the trial order was pseudorandomised such that stop or change trials did not occur more than twice in a row. The same colour counterbalancing was used in the SCT as in the SST.

### Data Analysis: Behavioural

Unless otherwise specified, statistical analyses were conducted using generalised linear mixed models (GLMMs) in Jamovi (The Jamovi Project, 2021). To run the GLMMs, the GAMLj module of Jamovi was used (Gallucci, 2019). A model selection approach based on Bayesian information Criteria (BIC; Schwartz, 1978) was used to determine the optimal structure of random effects.

Prior to analysing reaction time (RT), trials with RTs < 50ms were excluded, on the basis that these trials represent pre-emptive movements. To analyse reaction times differences in correct go trials, we used a GLMM with a gamma distribution and a log link function, which accommodates for the inherent positive skew in RT data (Lo and Andrews 2015). Fixed effects were condition (CRT, SST, SCT) and congruence (congruent, incongruent). The selected random effects structure included participant intercepts and slopes for both fixed effects. Notably, the difference in go RTs between the CRT and SST and SCT can provide an index of “proactive slowing”, that is, the degree to which participants slowed their movement enaction in anticipation of the stop/change signal (see Langford et al., 2016).

To test if congruence of the go response had an effect on the speed of the subsequent RT in correct change trials, an additional GLMM (with a gamma distribution and log link function) was run comparing “change time” (i.e., the response time minus the SCT for that trial, RT-SCT) for congruent and incongruent change trials. The fixed effects structure included the single factor of congruence, and the selected random effects structure included participant intercepts and slopes for congruence.

For both the SST and SCT, SSRT was calculated separately for congruent and incongruent trials using the integration method (Verbruggen et al., 2019). A linear mixed model (Gaussian distribution and identity link) with a random effects structure of participant intercepts was run on this data.

### Data Analysis: Electromyography (EMG)

EMG data were processed using MATLAB (MathWorks, 2018) following algorithms developed in our previous work and available online (Salomoni et al. 2025). Specifically, EMG signals were initially filtered by a fourth-order band-pass Butterworth filter at 20–500 Hz. Signals were rectified and subsequently low-pass filtered at 50 Hz. The detection of EMG onset and offset was achieved through a single-threshold algorithm (Hodges & Bui, 1996). A sliding window of 500ms was used to establish the lowest root mean squared amplitude within each trial segment, which then served as a baseline. Bursts were considered significant when the smoothed EMG exceeded three standard deviations (SDs) above baseline, with bursts less than 20ms apart merged into a single event.

Two types of EMG bursts were identified: the *RT-generating burst*, which occurred after the go stimulus but *before* the recorded button press, and *partial bursts*, which started decreasing before sufficient force was applied to elicit an overt button press. To identify partial bursts, the peak amplitude of the responding hand was required to exceed 10% of the average peak amplitude from the participant’s successful go trials in that condition. This process helped exclude activity unrelated to the task stimuli. For each burst, onset and offset times (relative to the relevant stimuli), as well as peak amplitude times, were extracted. The algorithm for identifying partial bursts avoided spurious inclusion of RT generating activity in the non-cued hand by combining bursts that are separated by less than 20ms. This ensured that only a full suppression of muscle activity followed by reactivation is recognized as a distinct partial burst.

A trial-level measure of stopping speed can be determined by subtracting SSD/CSD from the time (in milliseconds) of the peak of partial bursts (i.e., the point at which muscle activity begins to decrease; Raud et al., 2022). Here we refer to this measure as CancelTime (Jana et al., 2020). The use of CancelTime as a valid measure of inhibition latency depends on a having sufficient number of trials with partial bursts. Notably, CancelTime is ‘censored’ at each end of its distribution: ‘late’ inhibition invariably results in failed stops or changes, and ‘early’ inhibition generally results in successful trials *without* partial bursts (Salomoni et al., 2025). To determine if congruence or stop/change conditions influenced the likelihood of occurrence of partial bursts, we ran a GLMM with a probit categorical dependent variable (the presence/absence of partial bursts) with factors of congruence (congruent, incongruent) and condition (SST, SCT). The selected random effects structure included participant intercepts and slopes for condition.

To analyse Canceltime a GLMM with a gamma distribution and log link function was used. We included fixed effects of congruence (congruent, incongruent) and condition (SST, SCT). The selected model included a random effects structure of participant intercepts and slopes for condition. To investigate the strength of evidence following a null-effect of congruence, Bayesian independent-samples t-tests were run on log-transformed Canceltimes (seperately for stop and change conditions), using a Cauchy prior distribution with a scale factor of 0.707 (Wagenmakers et al., 2018).

We also used EMG to attain a trial-level index of the speed of action reprogramming following a successful stop with a partial burst in the SCT, to determine if this was influenced by congruency. Action reprogramming was calculated as the time from peak of the partial burst to the onset of the RT-generating burst. The model included the single fixed factor of congruence (congruent, incongruent). Following visual inspection of distributions, the positive skew on this data was minimal compared to RT data. Here, we compared BICs across both Gaussian distributions with Identity links and Gamma distributions with log links. The final random effects structure used Gaussian distributions and identity link, with participant intercepts in the random effects structure.

## Results – Experiment 1

### Behavioural Results

Response rates and accuracy in go trials suggested that participants attended to the task well, with few missed attempts. Success rates were close to 50% in the stop and change trials of the SST/CST for both congruent and incongruent trials, indicating the SSD/CSD staircasing procedure was effective. See Table 1 for a summary of key behavioural results.

**Table 1.** Summary of Key Behavioural Results in Experiment 1.

| Condition | Congruence | Correct go RT (ms) | Correct Change RT (ms) * | Go % Correct | Mean SSD/ CSD (ms) | Stop/Change % Correct |
| --- | --- | --- | --- | --- | --- | --- |
| **CRT** | Congruent | 375 [338-415] |  | 99.41% |  |  |
|  | Incongruent | 418 [379-461] |  | 95.82% |  |  |
| **SST** | Congruent | 486 [450-525] |  | 99.54% | 204 | 49.34% |
|  | Incongruent | 517 [480-558] |  | 99.50% | 223 | 51.07% |
| **SCT** | Congruent | 474 [432-521] | 426 [401-452] * | 99.22% | 241 | 50.60% |
|  | Incongruent | 503 [460-550] | 427 [401-455] * | 98.87% | 256 | 52.16% |

RT values represent estimated marginal means from associated models [95% CIs in brackets]. All other values are descriptive data. \*These values represent time from presentation of the change signal (at CSD) until the correct reaction time.

#### Reaction Time (RT)

The results for analysis conducted on go RTs are depicted in Figure 2. The model revealed significant main effects of congruence, χ^2^(1) = 18.277, *p* <.001, condition, χ^2^(2) = 34.380, *p* <.001, and a significant interaction, χ^2^(2) = 8.880, *p* =.012. Bonferroni corrected post-hoc tests revealed significant effects of congruence in all conditions (CRT: *z* = 4.63, *p* <.001; SST: *z* = 3.56, *p* =.006; SCT: *z* = 3.36, *p* =.011) with faster responses in congruent relative to incongruent trials. Pairwise post-hoc tests on condition revealed faster RTs in the CRT condition compared to both the SST, *z* = 5.867, *p <*.001, and the SCT, *z* = 4.616, *p* <.001, but no significant difference in RTs between the SST and SCT, *z* = 0.947, *p* =.267.

**Figure 2:**
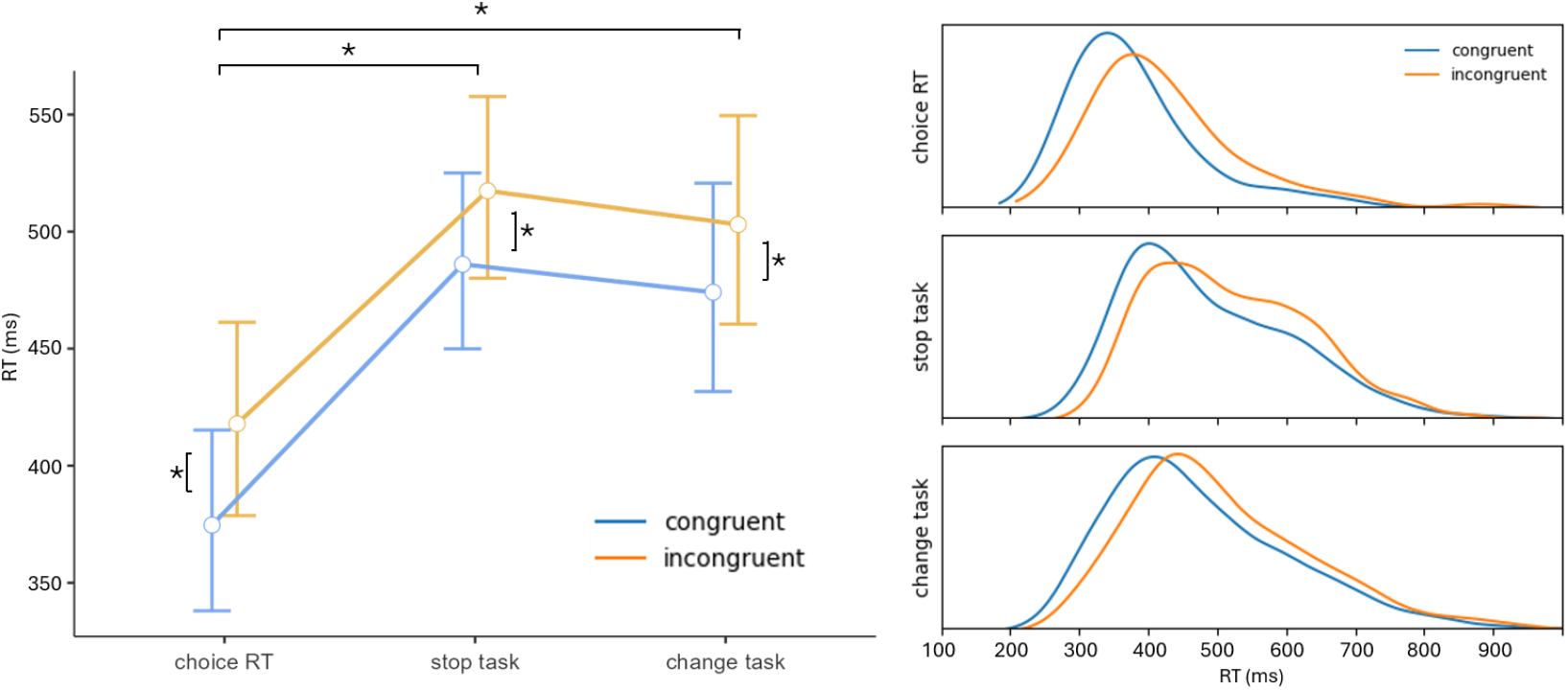
Correct go trial reaction times for congruent and incongruent trials in choice response, stop signal, and stop change tasks in Experiment 1. The slower reaction times and greater positive skew in the stop and change tasks is a result of proactive slowing, in anticipation of the need to stop. Error bars represent 95%CIs. * = *p* < 0.001

#### Change Reaction Time

The analysis on change reaction times revealed no significant main effect of congruence χ^2^(1) = 0.065, *p* = 0.799. Estimated marginal means are presented in Table 1.

#### Stop Signal Reaction Time (SSRT)

The race model, upon which SSRT calculations are based, requires that mean RT in failed stop or change trials is faster than mean go RT. This assumption was met by all participants in all conditions. The model conducted on SSRT revealed significantly longer SSRTs in the SST compared to the SCT, *F*(1) = 88.49, *p* <.001, no significant effect of congruence, *F*(1) = 0.800, *p* =.374, and no significant interaction effect *F*(1) = 0.135, *p =* 0.714. Estimated marginal means are presented in Table 2.

**Table 2.**
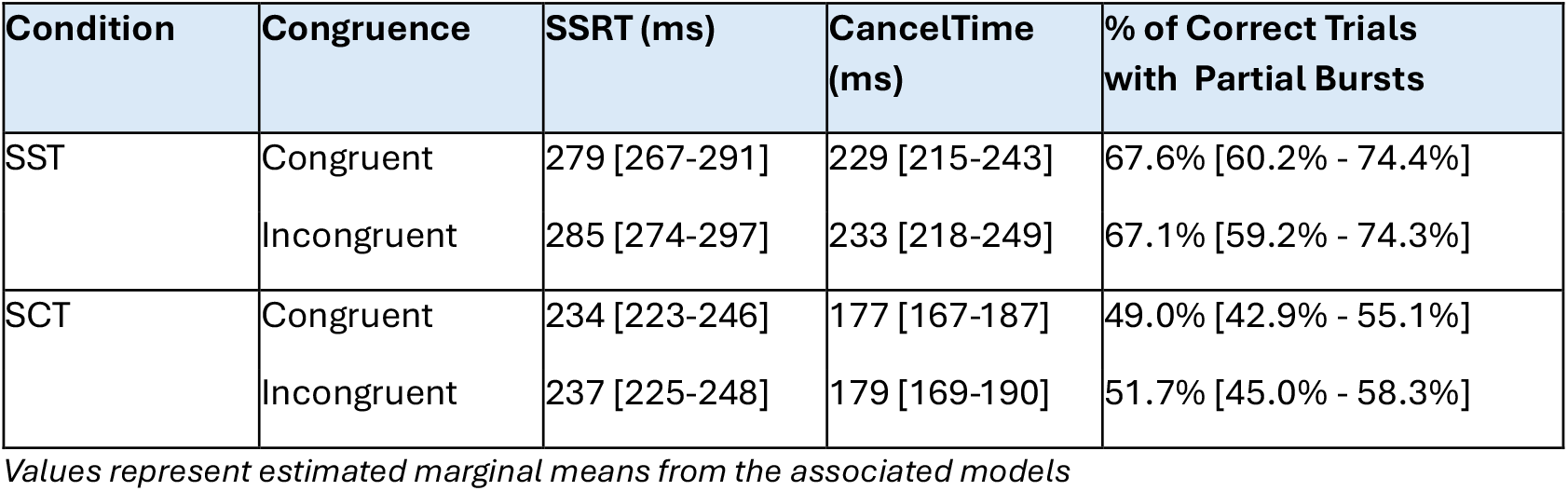
Behavioural and Physiological Indices of Stopping in Experiment 1.

### Electromyography (EMG) Results

#### CancelTime

Results from the model run on CancelTime are depicted in Figure 3. The GLMM on CancelTime revealed significantly slower stopping in the SST compared to the SCT, χ^2^(1) = 103.305, *p* <.001, no effect of congruence, χ^2^(1) = 2.177, *p* = 0.140, and no significant interaction, χ^2^(1) = 0.087, *p* =.769, corroborating the results of the SSRT analysis. Table 2 shows EMMs of CancelTime values for congruent and incongruent trials. Follow-up independent samples Bayesian t-tests confirmed strong evidence for no effect of congruence in the change task BF01 =13.64 and moderate-strong evidence for no effect in the stop task BF01 = 7.72.

**Figure 3:**
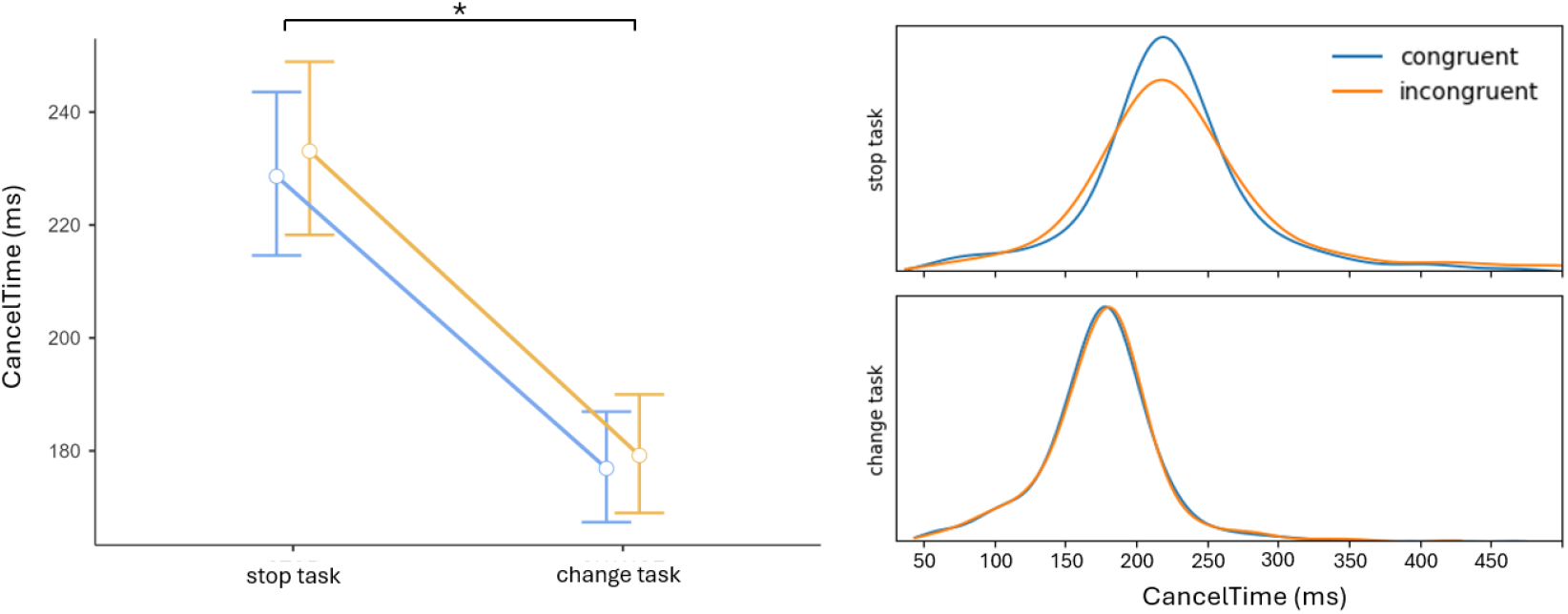
CancelTimes for congruent and incongruent trials in stop signal and stop change tasks in Experiment 1. Error bars represent 95%CIs. * = *p* < 0.001

#### Frequency of Partial Bursts

The model revealed no significant effect of congruence, χ^2^(1) = 0.419, *p* = 0.518, however there was a significant effect of condition, χ^2^(1) = 19.019, *p* <.001, whereby a significantly greater number of partial bursts were detected in the stop condition (refer to table 2). No significant interaction was observed χ^2^(1) = 1.063, *p* =.303.

#### Action Reprogramming

Distributions of action reprogramming times from the change task (and a schematic explaining how these were calculated, and how they relate to CancelTime) can be found in Figure 4. The model revealed a significant effect of congruence, *F*(1) = 5.887, *p* = 0.015, whereby action reprogramming was faster in congruent trials, 350ms 95%CIs[305-396ms], than incongruent trials, 364ms 95%CIs[318-410ms].

**Figure 4.**
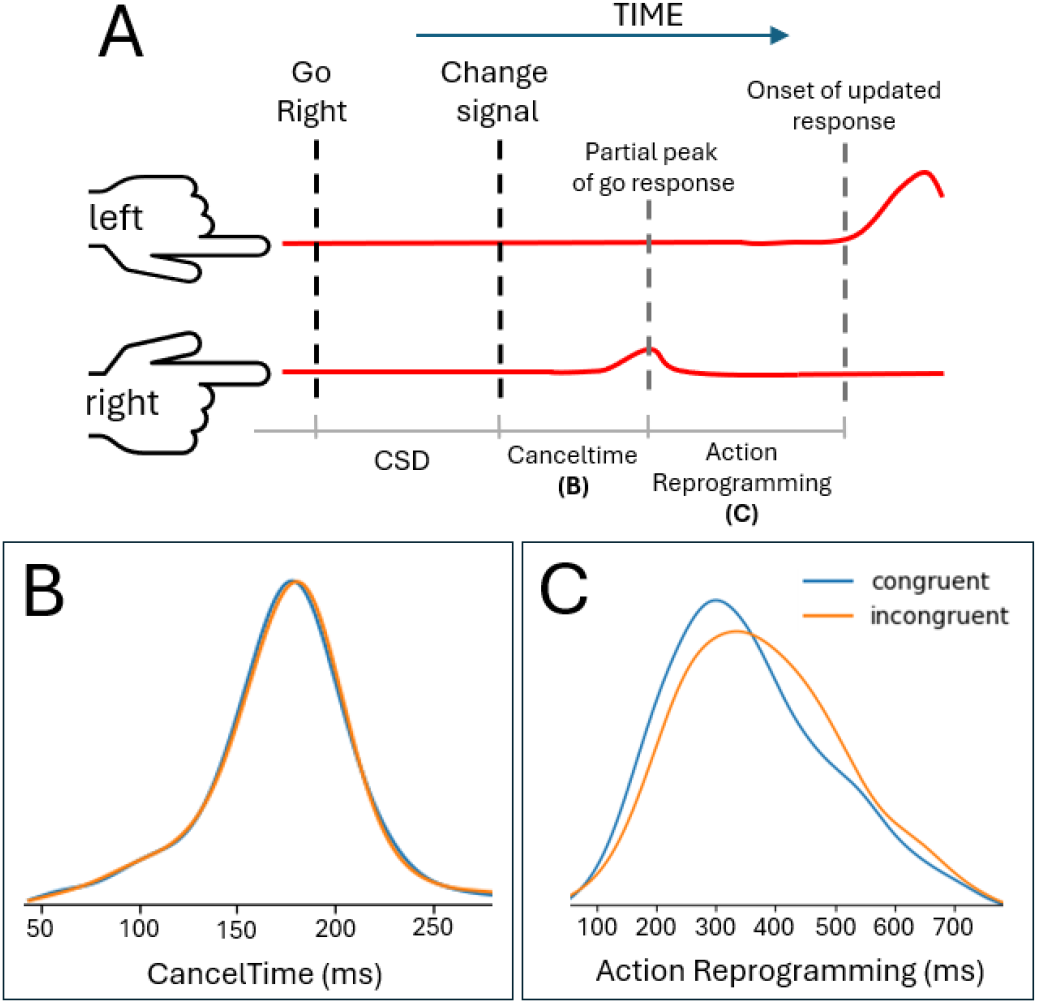
**A)** schematic representing hypothetical smoothed EMG amplitude during a change trial that started with a right go. CancelTime represents the time between the appearance of the change stimulus (following the change signal delay; CSD) and the peak of the partial response. The time from the peak of the partial response to the onset of the correct response (cued by the change signal) represents action reprogramming time. **B)** CancelTime distributions demonstrated no significant differences between congruent and incongruent trials **C)** Distributions of action reprogramming, which was significantly faster in congruent trials.

**Figure 5:**
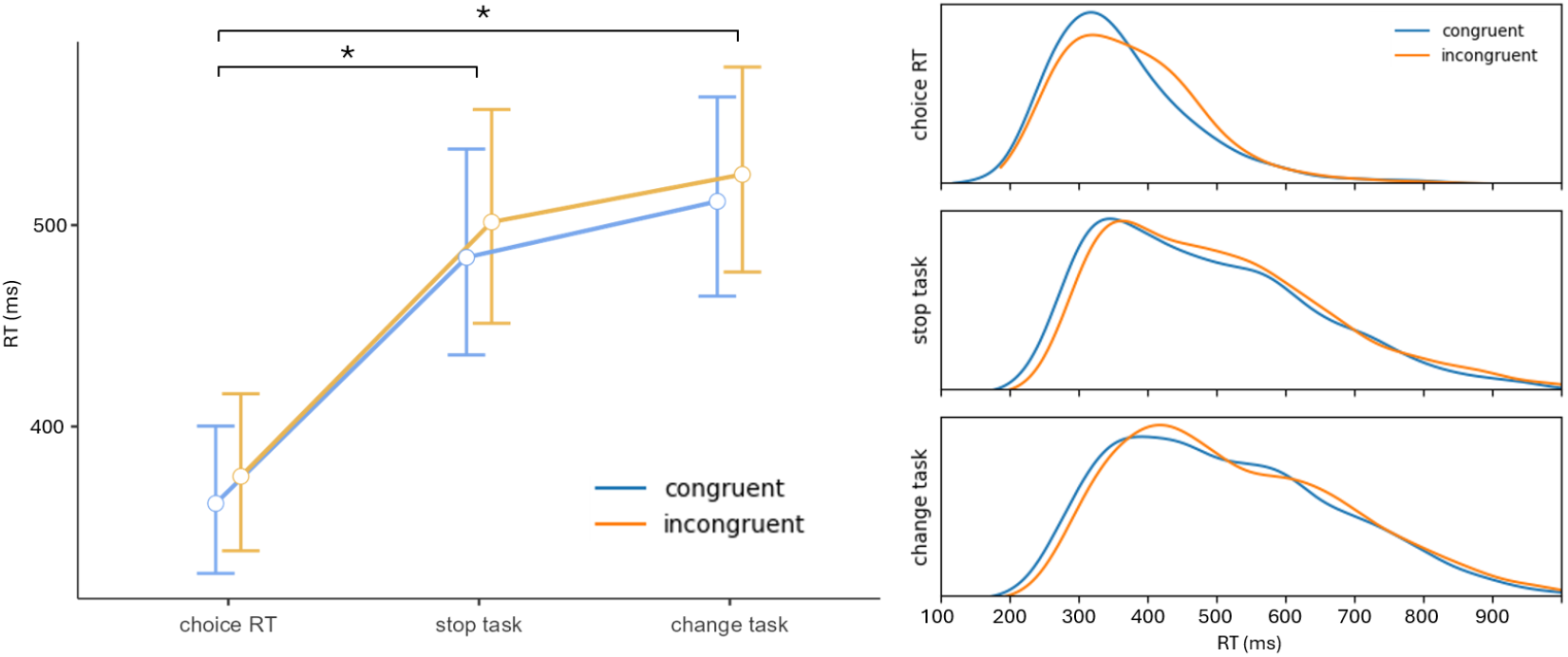
Correct go trial reaction times for congruent and incongruent trials in choice response, stop signal, and stop change tasks in Experiment 2. There was also a significant effect of congruence, though no interaction with condition, so this was not followed up with separate assessments of congruence in each condition. Error bars represent 95%CIs. * = *p* < 0.001

## Experiment 2

Following the unexpected observation of faster SSRT and CancelTime in the change condition compared to the stop condition, we ran a follow-up experiment to control for possible stimulus effects. In Experiment 1, the stop stimulus changed *colour* whereas the change stimulus changed *location* (see Figure 1). While similar use of stimuli has been used in other stop change tasks (e.g., Hervault & Wessell, 2025), there is some evidence that when stop signals are presented at a different location to the go signal, stopping speed is hastened relative to when a stop signal that is superimposed over the go signal (Friehs et al., 2024). As such, we ran a second experiment with a new cohort of participants, in which both stop and change trials featured a colour change (c.f. the stop trials from Experiment 1). In all other ways, the experiment design and procedure were identical to that of Experiment 1. While a distinct design could have been used to specifically address this question (e.g., a within subject experiment specifically testing stimulus effects in SCTs), we opted to use the opportunity to determine replicability of other key results (e.g., congruency effects on go and stop responses) from Experiment 1, to ensure that the same conclusions could be drawn in a distinct (albeit, demographically similar) cohort (Nosek et al., 2022).

### Participants

Twenty-five healthy participants (16 female; M_Age_ = 27.8 years, SD = 7.5 years) were recruited via the same methods as those described for Experiment 1. As per Experiment 1, both stimulus colour and task order were counterbalanced.

### Analyses

Each of the analyses described in Experiment 1 was also conducted on the data from Experiment 2. While the same fixed effects structures were used in Experiment 2, model selection (based on BIC) for random effects structures was run independently for this cohort. Ultimately, this led to the same random effects structures as described in Experiment 1 for all analyses, barring the reaction time analysis on correct go trials, for which the random effects structure included participant intercepts and slopes for condition. As in Experiment 1, the strength of evidence for null effects with regards to Cancel Times was assessed with follow-up Bayesian independent samples t-tests.

## Results

The key behavioural results for experiment 2 are presented in Table 3.

**Table 3.** Summary of Key Behavioural Results in Experiment 2.

| Condition | Congruence | Correct go RT (ms) | Correct Change RT (ms) * | Go % Correct | Mean SSD/CSD (ms) | Stop/Change % Correct |
| --- | --- | --- | --- | --- | --- | --- |
| **CRT** | Congruent | 362[327-400] |  | 99.8% |  |  |
|  | Incongruent | 375[338-416] |  | 97.3% |  |  |
| **SST** | Congruent | 484[436-538] |  | 98.4% | 253 | 50.3% |
|  | Incongruent | 502[451-557] |  | 98.5% | 269 | 51.3% |
| **SCT** | Congruent | 512[465-564] | 433 [396-474]* | 97.3% | 276 | 51.5% |
|  | Incongruent | 525[477-578] | 438 [395-485]* | 95.8% | 291 | 53.4% |

RT values represent estimated marginal means from associated models [95% CIs in brackets]. All other values are descriptive data. \*These values represent time from presentation of the change signal (at CSD) until the correct reaction time.

### Reaction Time (RT)

The model run on reaction time data revealed significant main effects of congruence, χ^2^(1) = 22.584, *p* <.001, condition, χ^2^(2) = 38.779, *p* <.001, but a non-significant interaction, χ^2^(2) = 1.832, *p* = 0.400. Pairwise post-hoc tests on condition revealed faster RTs in the go-only condition compared to both the SST, z = 5.710, *p <*.001, and the SCT, z = 6.153, *p* <.001, but no significant difference in RTs between the SST and SCT, z = 1.701, p = 0.267.

### Change Reaction Time

The analysis on change reaction times revealed no significant main effect of congruence χ^2^(1) = 0.393, *p* = 0.530. Estimated marginal means are presented in Table 3.

### Stop Signal Reaction Time (SSRT)

One participant was removed from SSRT calculations in Experiment 2 on account of violations of the race model that underpins its calculation (mean failed stop RT was not faster than go RT in two conditions). The model run on the remaining participants revealed no significant main effect of condition *F*(1) = 0.098, *p* = 0.756, no significant effect of congruence, *F*(1) = 0.530, *p* = 0.469, and no significant interaction *F*(1) = 0.070, *p* = 0.792. Estimated marginal means are presented in Table 4.

**Table 4.** Behavioural and Physiological Indices of Stopping in Experiment 2.

| Condition | Congruence | SSRT (ms) | CancelTime (ms) | % of Correct Trials with Partial Bursts |
| --- | --- | --- | --- | --- |
| SST | Congruent | 229 [210-249] | 155 [144-166] | 70.7% [64.8% - 76.1%] |
|  | Incongruent | 226 [207-246] | 155 [144-167] | 70.1% [63.6% - 75.9%] |
| SCT | Congruent | 233 [214-253] | 158 [146-170] | 57.2% [52.0% - 62.2%] |
|  | Incongruent | 227 [207-246] | 163 [150-177] | 60.5% [54.8% - 66.0%] |

*Values represent estimated marginal means from the associated models*

### Electromyography (EMG) Results

#### CancelTime

Results from the Experiment 2 CancelTime analysis are depicted in Figure 6. In contrast to Experiment 1, the GLMM on CancelTime revealed no significant effects of condition, *χ*^*2*^(1) = 1.473, *p* = 0.225, or congruence, *χ*^2^(1) = 1.408, *p* = 0.235, and no significant interaction, *χ*^*2*^(1) = 1.319, *p* =.251. The follow-up Bayesian t-tests revealed strong evidence for the null with regards to congruence BF01 = 12.05, but only anecdotal evidence for the null with regards to condition BF01 = 1.69. Table 4 shows EMMs of CancelTime values for congruent and incongruent trials.

**Figure 6.**
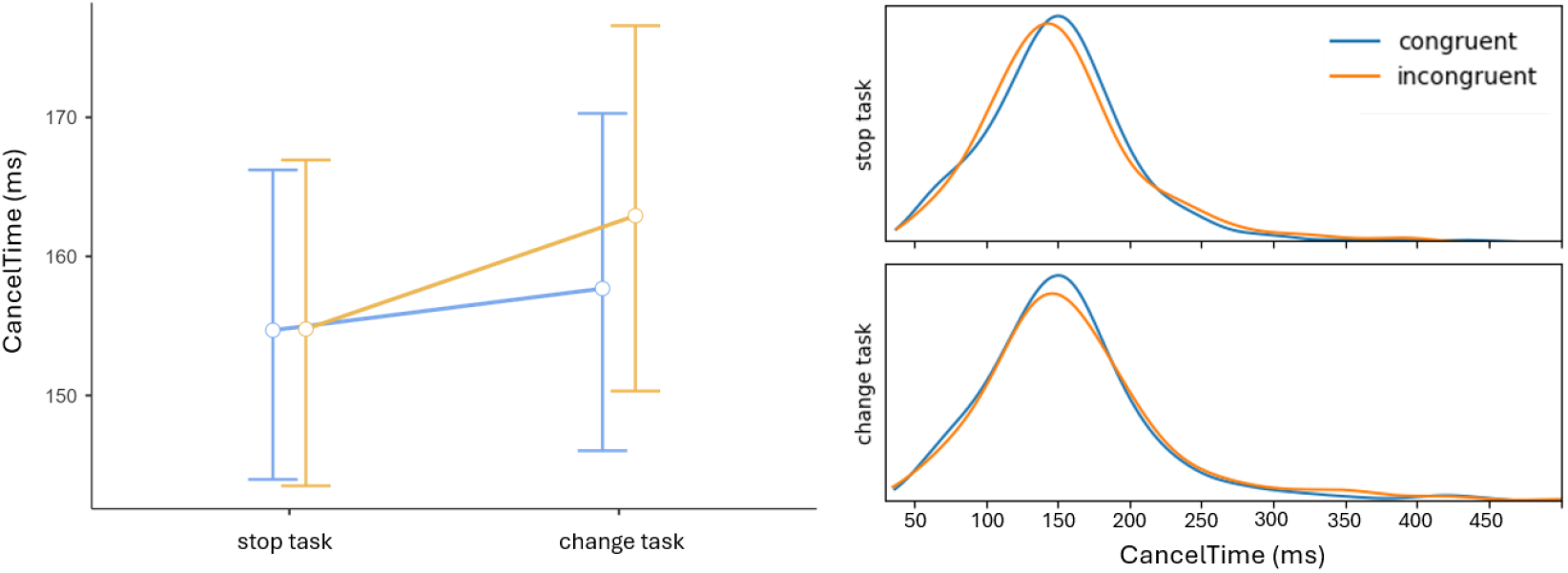
CancelTimes for congruent and incongruent trials in stop signal and stop change tasks in Experiment 2. Error bars represent 95%CIs. * = *p* < 0.001

#### Frequency of Partial Bursts

The model revealed no significant effect of congruence, χ^2^(1) = 0.677, *p* = 0.410, but there was a significant effect of condition, χ^2^(1) = 20.380, *p* <.001, whereby a significantly greater number of partial bursts were detected in the stop condition (refer to table 2). No significant interaction was observed χ^2^(1) = 1.654, *p* =.198.

#### Action Reprogramming

The model revealed a significant effect of congruence, *F*(1) = 11.209, *p* < 0.001, whereby action reprogramming in the stop change condition was faster in congruent trials 411ms 95%CIs[354-468ms], than incongruent trials 431ms 95%CIs[374-489ms].

## Discussion

The current study used novel versions of stop signal tasks (SSTs) and stop change tasks (SCTs) to assess whether biasing a movement to be expected or unexpected influenced the speed with which it could be cancelled or adapted. As expected, biasing information effectively influenced the speed of action execution; however, this biasing did not influence the speed at which that action could be subsequently cancelled. Physiological results also indicated that expected movements can be reactively reprogrammed more quickly than unexpected movements. Interestingly, a second experiment replicated these results and additionally suggests that the stop process is sensitive to stimulus features (specifically, that stopping is faster when the stop signal is presented in a separate location to the go signal). When stimulus effects were controlled for, there was no difference in stopping speeds between the SST and SCT. These findings add to the body of evidence suggesting mechanisms in the SCT are the same as those in the SST, with the addition of a new action (Boecker et al., 2011). This also aligns with contemporary models of stopping where the stop process is independent of any subsequent actions (Gronau et al., 2024, Salomoni et al., 2025).

### Priming a movement does not significantly influence the speed of its subsequent cancellation

The current experiments corroborate findings that probabilistic cues informing participants of the likelihood that a particular movement will be required significantly influence RTs (Bestman et al., 2008; Garton et al., 2019; Miller & Low, 2001). Our follow-up Bayesian tests provide strong evidence that, despite this influence of movement enaction, the subsequent cancellation of that movement remains unaffected. Conventional stop signal tasks represent highly artificial environments, and in day-to-day scenarios, reactive action cancellation occurs in the context of ongoing motor plans, enacted based on prior knowledge and expectations. A key challenge in inhibition literature is that it is currently unclear how well stopping tasks relate to the real world (Hannah & Aron, 2021). Establishing what does, and does not, influence stopping mechanisms represents one of the main means by which this challenge can be addressed. The current experiments contribute to this understanding by demonstrating that reactive stopping either overrides, or operates in parallel with, expectations that bias the movements we plan and initiate.

The finding that biasing the go process does not influence the speed of stopping raises a number of questions regarding the neural mechanisms that underpin action planning, enaction and cancellation. While the mechanisms by which prior expectation influence RT are incompletely understood, brain stimulation research has identified specific motor-system adjustments following biasing cues like those used in the current tasks. For instance, during movement preparation in choice response tasks, there is a general suppression of corticospinal excitability (Duque et al., 2017), and further effector-specific suppression if biasing information renders a particular movement more likely (Bestmann et al., 2008). This coincides with local *increases* in M1 excitability for that effector (Tandonnet et al 2010). The combination of local excitation along with general corticospinal suppression may facilitate movement preparation while preventing premature release of either action choice (Duque et al., 2017). In terms of how this relates to the manifest behaviour: the increased facilitation for the cued hand purportedly results in less requisite neural activation upon presentation of the go stimulus, enabling faster responses for congruent trials. At the same time, the hand representation that is *less* likely to require a response is subject to a greater amount of inhibitory signalling from the opposite hemisphere (Puri & Hinder, 2022). As such, the non-cued hand likely requires greater recruitment of motor neurons and/or greater firing rates for a response to be generated, manifesting behaviourally as *slower* response times for incongruent trials (Puri et al., 2023). While the pathways underpinning reactive action cancellation also culminate in M1 (Aron, 2011; Diesberg and Wessell 2021), the current result suggests reactive inhibitory mechanisms must either immediately override these preparatory modulations, or operate in parallel. The way in which the motor system accommodates this, despite both mechanisms converging on M1, is yet to be explained. Previous research observed that when cues yielded information about which components of a *bimanual* go response may require cancellation, it was the speed of the ongoing action, not the speed of the stopping that was affected (Salomoni et al., 2023). Collectively, this suggests that preparatory changes that accompany expected movements influence the enaction of movement both prior to, and after stopping, but the stop mechanism itself, remains independent of these preparations.

Future research should test the generalisability of this finding. Here, we specifically used trial-level probability information, but further research may seek to compare block-level biasing information (e.g., “70% left” for all trials in a block) to more flexible trial-wise biasing (Garton et al. 2019), or use other established sources of go response bias, such as greater reward on one side (Bundt et al., 2019; Klein et al., 2012). To our knowledge, the only current paper to investigate this effect did so in an implicit manner: Rather than explicitly providing bias information to participants, the experiment manipulated the probability of left and right responses (e.g., 30% of trials left go, 70% trials right go) and allowed participants to learn this information over time (i.e., this probability information was not explicitly told to participants; Federico & Mirabella, 2014). Despite finding no main effect of probability on RTs, SSRTs were significantly *faster* for responses required on the side that was less likely to require movement. This study involved reaching movements, rather than ballistic button-press movements, and only used behavioural estimates of stopping, likely explaining the difference in results. Nonetheless, it highlights that further research using larger-scale limb movements is needed to establish the contextual specificity of the current result.

### Expected movements can be reprogrammed more quickly than unexpected movements

In both experiments, we observed that action reprogramming (the time between the suppression of the initial response and initiation of the updated response in the contralateral effector) was faster for congruent than incongruent trials. Notably, in incongruent change trials, the participant has just been subject to two infrequent stimuli in rapid succession (a go cue on the unexpected side, and then a change cue – which also occurs only infrequently). Unexpected, or infrequent events are known to trigger a transient interruption to motor output (Wessel & Aron 2017), delaying any ongoing or subsequent responses. This is thought to be underpinned by the hyperdirect pathway, connecting inferior frontal cortices with basal-ganglia motor outputs and rapidly interrupting output to M1 (Diesberg & Wessel, 2021). Upon being triggered, this mechanism is thought to facilitate the retuning of subsequent movement (Hervault and Wessel, 2025). It is currently unclear a) what is sufficient to trigger this mechanism and b) how regularly it can be triggered. A possible explanation for the current finding is that processing and responding to unexpected stimuli depletes a common resource, resulting in slower adaptions when numerous unexpected events occur in rapid succession. If such a mechanism can only be triggered infrequently, and facilitates the rapid updating of motor plans, this could explain the slower retuning of movements that were already rendered unexpected by biasing cues. Alternatively, it may be that, while the latency of stopping wasn’t affected, the *degree* of inhibition that occurs to suppress unexpected movements is greater, requiring more excitation to be overcome, and leading to slower muscle recruitment of the subsequent movement. Future research could provide further insights by using transcranial magnetic stimulation to assess the excitability of the corticospinal tract (with motor evoked potentials; Bestman & Krakauer, 2015) following the suppression of expected and unexpected movements. MEPs could also be used to determine if the degree of transient suppression of corticospinal excitability that is known to occur following unexpected stimuli (Iacullo et al., 2020) sequentially reduces if numerous distinct unexpected events occur in a row. To the author’s knowledge, this remains untested.

### The speed of the stopping process is highly sensitive to stimulus effects

The results of Experiment 2, where stimuli for stop and change conditions were identical, align with some previous work demonstrating that stopping speed does not significantly differ between SSTs and SCTs (Boecker 2011; Drueke et al., 2010). This suggests that the fast, automatic stopping process is unaffected by whether stopping occurs in isolation, or is followed by the requirement to implement an altered or new response. Moreover, this finding is consistent with a recent model of complex stopping where a global stop process occurs independently to action reprogramming of movement components that are still required to be enacted following a stop signal (Gronau et al., 2024, Salomoni et al., 2025). Neuroimaging research contrasting stop and change trials supports the notion that the same stopping mechanism is implemented for both contexts, based on the fact that significant differences between the two can be wholly explained by the additional requirements of enacting a second (changed) response. Specifically, in change trials (contrasted with stop trials) there is greater activity in M1, premotor cortices, postcentral gyrus, and the inferior parietal lobe. Notably, no differences in activity is observed in regions typically associated with action cancellation (e.g., right inferior frontal gyrus, pre-supplementary motor area; Boecker 2011). Notably though, these findings (and those of Drueke et al., 2010) were from a task which intermixed change and stop trials within the one condition. Other research implementing SSTs and SCTs as *distinct* tasks have instead observed faster SSRTs in SSTs than SCTs (De Jong et al., 1995; McClure et al., 2005). Some authors have speculated that this difference may be due to greater task difficulty and working memory demands in SCTs compared to SSTs (and that these demands would not be specific to change trials if both stopping and changing are integrated into the one, more complex condition; Boecker et al., 2013). The reason that we didn’t observe a difference between SSTs and SCTs (aside from those arising from stimulus effects once stimulus effects in Experiment 1), despite using a block-wise design, is not immediately clear.

The absence of an observable difference in stopping speed between the SST and SCT in Experiment 2 also contrasts with a recent paper by Hervault and Wessell (2025) which reported *faster* stopping (by both SSRT and EMG-measures) in an SCT relative to an SST. Notably though, we observed a similar effect in Experiment 1 of the current study. Both Experiment 1, and the study by Hervault and Wessell (2025), signalled stop change trials with an additional stimulus appearing. When we replicated the experiment without the more salient stimulus in the SCT (i.e., using only a colour change for both SST and the SCT), stopping speed was no longer significantly different between these tasks. This corroborates other resent research which observed that presentation of the stop signal in a distinct location to the go signal significantly shortens SSRTs (Friehs et al., 2024). Notably, this stimulus effect is rarely considered in SST task design (Verbruggen et al., 2019), as both colour-change (e.g., Cai & Leung, 2009; Ganos et al., 2014; Jana et al., 2020; Weber et al., 2025), and additional (distinct) stimulus stop signal variants (e.g., Sharp et al, 2010; Schmitt et al., 2017) are common. More salient stop signals have been associated with shorter SSRTs, irrespective of location (Kenemans et al., 2023). In the SCT, there were two salient changes (the initial stimulus disappearing, and a distinct stimulus appearing), while in the SST there was only one (the colour change). A comparison of stopping speeds in a SSTs and SCTs using auditory stop signals (that keep stimuli consistent but vary only instructions that precede the task) would be a logical extension to confirm the generalisability of current findings (Weber et al., 2024b).

### Key results remained robust, but replication resulted in notable differences

Running a replication of Experiment 1 (baring the modified stimulus in the SCT) provided an opportunity to assess the robustness of key findings with regard to how bias affects response execution, cancellation and subsequent changing, while also allowing us to assess the effects of stimulus features on stopping/change performance. Perhaps the most notable unexpected difference between the two experiments was the overall slower stopping speeds observed in Experiment 1 than Experiment 2, especially in the SST. Furthermore, congruency effects were qualitatively smaller in Experiment 2. Given the same protocol and devices were used for both experiments, the only plausible explanation for this difference is cohort effects. While participants were recruited from the same sources, the data was collected at different times of year, and there may have been systematic differences that can only be speculated at, but which apply to the majority of behavioural experiments. This replication demonstrates the robustness of our key results, but we note that these differences also highlight the importance of prioritising single-session within-subject comparisons for inhibition research, whenever possible (Thunberg, 2024).

### Conclusion

A key challenge in inhibition research is that conventional experimental paradigms represent a highly artificial scenario, and it is unclear how well these generalise to the real world. The current experiments contribute to our broader understanding of this relationship by demonstrating that reactive stopping mechanisms are utilised consistently for both expected and unexpected movements, and invariant to whether the stop is followed by another action. We also provide evidence that the reprogramming of movement can be performed more quickly for expected than unexpected movements. The neural mechanisms for this observation are currently uncertain, highlighting a potential avenue for future research.

## Credit Statement

Simon Weber – Conceptualisation, Methodology, Formal Analysis, Software, Writing - Original Draft Preparation

Nicholas Vukac - Conceptualisation, Investigation, Methodology, Formal Analysis, Writing - Original Draft Preparation

Sauro E. Salomoni – Conceptualisation, Formal Analysis, Methodology, Software

Alison J. Ross - Conceptualisation, Investigation, Data Curation

Emily Coleman - Conceptualisation, Investigation, Data Curation

Mark R. Hinder - Conceptualisation, Supervision, Writing - Review & Editing

The data associated with this manuscript can be found on OSF https://osf.io/dzm4f/

Our MATLAB scripts for analysing EMG data can also be found on OSF: https://osf.io/r8a54

